# Distributed Feedforward and Feedback Processing across Perisylvian Cortex Supports Human Speech

**DOI:** 10.1101/2021.12.06.471521

**Authors:** Ran Wang, Xupeng Chen, Amirhossein Khalilian-Gourtani, Leyao Yu, Patricia Dugan, Daniel Friedman, Werner Doyle, Orrin Devinsky, Yao Wang, Adeen Flinker

## Abstract

Speech production is a complex human function requiring continuous feedforward commands together with reafferent feedback processing. These processes are carried out by distinct frontal and posterior cortical networks, but the degree and timing of their recruitment and dynamics remain unknown. We present a novel deep learning architecture that translates neural signals recorded directly from cortex to an interpretable representational space that can reconstruct speech. We leverage state-of-the-art learnt decoding networks to disentangle feedforward vs. feedback processing. Unlike prevailing models, we find a mixed cortical architecture in which frontal and temporal networks each process both feedforward and feedback information in tandem. We elucidate the timing of feedforward and feedback related processing by quantifying the derived receptive fields. Our approach provides evidence for a surprisingly mixed cortical architecture of speech circuitry together with decoding advances that have important implications for neural prosthetics.

## 1 INTRODUCTION

The central sulcus divides the human frontal from the posterior temporal, parietal, and occipital neocortices [34]. Traditionally, this divide separates high order planning and motor execution from sensation. Feedforward execution lies in the frontal cortices in contrast to feedback sensory processing across posterior cortices for the various sensory modalities (e.g., auditory, visual, somatosensory, etc.) [17]. Higher order capacities such as working memory, cognitive control, and decision making are often viewed as initiated by frontal cortices with direct influence on sensory cortices [19, 38, 44].

Human higher order cognitive functions include planning and executing complex speech sequences that carry semantic and linguistic meaning [7,29]. Speech production is a complex human motor behavior requiring precise coordination of multiple oral, laryngeal and respiratory muscles [42]. These finely tuned motor actions then produce reafferent feedback in the auditory, tactile, and proprioceptive domains as we process our own speech.

The dynamic influence of feedforward commands on sensory feedback is a hallmark of sensory motor systems across the animal kingdom [10]. For example, motor neurons in cricket both drive the generation of chirping sounds as well as inhibit the auditory system to filter out the loud noise produced by its wings [49]. Similarly, auditory neurons in the marmoset monkey are suppressed during vocalization to provide increased sensitivity to vocal feedback [50]. Prevailing models in human speech motor control propose a feedforward system that predicts and generates actions and a feedback system responding to the vocal auditory and somatosensory effects [22, 23, 26–28, 30]. There is a consensus that the two systems are anatomically separated, with the feedforward system mainly supported by ventral frontal cortices, while posterior cortices support feedback processing. During feedback processing, frontal cortices need to update new actions dynamically, and it remains unclear which subregions are involved in this process. Moreover, the exact timing of feedforward and feedback engagement across the cortex remains unknown.

A growing literature has leveraged unique human electrocorticographic (ECoG) recordings from patients undergoing neurosurgical procedures to obtain a combined spatial and temporal resolution critical for investigating speech production. Studies have detailed the signatures of feedforward speech planning [16] and organization of execution [5, 8] in frontal cortices as well as the subsequent auditory feedback architecture in temporal cortices [15,20,21]. To date, evidence of feedback processing has mainly focused on artificially altering the acoustic feedback to create a mismatch in the perceived pitch, providing evidence for enhanced responses in posterior cortices as well as frontal cortex (i.e., ventral sensorimotor cortex) [6]. However, the acoustic perturbation also causes speakers to compensate and change their produced pitch, leading to motor enhancement confounded with the feedback. Recently, the unprecedented signal-to-noise ratio offered by ECoG recordings has ushered deep neural network approaches to decode speech represented in auditory accurately [1,3,46,47] and sensorimotor [4] cortices. Nevertheless, these approaches have not been able to disentangle feed-forward and feedback contributions during speech production as the motor and sensory responses co-occur.

We directly disentangle feedback and feedforward processing during speech production by applying a novel deep learning architecture on human neurosurgical recordings to decode speech (Figure 1). Our approach decodes interpretable speech parameters from cortical signals, which drives a rule-based differentiable speech synthesizer. By learning neural network architectures which apply either casual (predicting using only the past), anticausal (predicting using the future feedback), or both (noncausal), spatial-temporal convolutions, we are able to analyze the overall feedforward and feedback contributions, respectively, as well as to elucidate the temporal receptive fields of recruited cortical regions. In contrast to current models that separate feedback and feedforward cortical networks, our analyses reveal a surprisingly mixed architecture of feedback and feedforward processing both in frontal and temporal cortices while achieving speech decoding performance on-par or better than previously reported.

**Figure 1:**
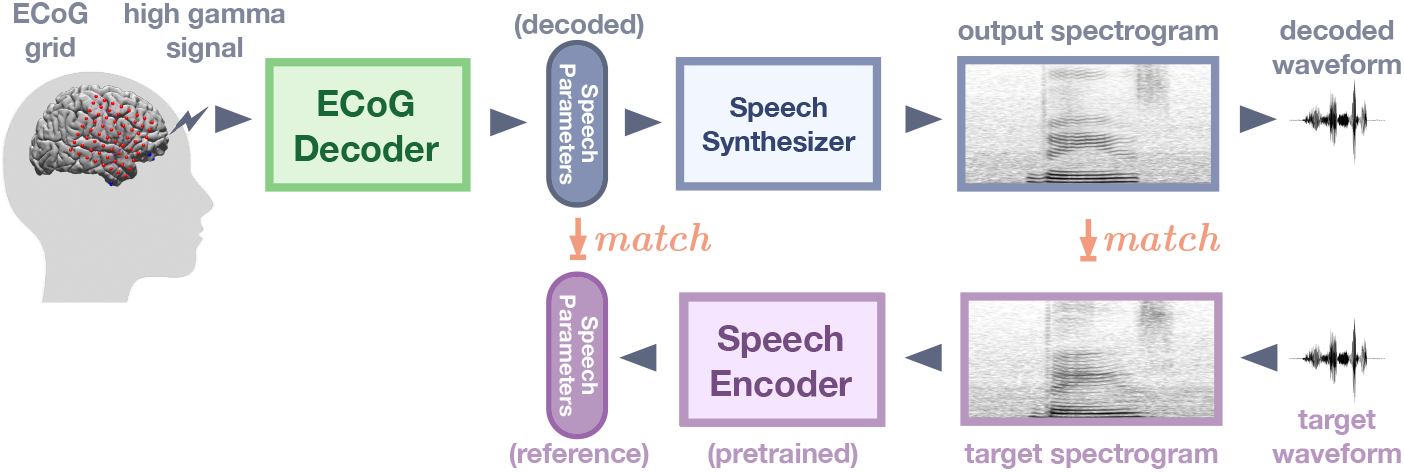
The overall structure of the decoding pipeline. ECoG amplitude signals are extracted in the high gamma range (i.e., 70-150 Hz). The ECoG Decoder translates neural signals from the electrode array to a set of speech parameters. This parameter representation is used to drive a speech synthesizer which creates a spectrogram (and associated waveform). During the training of the ECoG decoder, the speech parameters are matched to a reference derived by a speech encoder pre-trained using an unsupervised approach (without costly manual annotations). This approach constrains the learnt speech parameters and provides naturalistic decoded speeches.

## 2 RESULTS

We report speech decoding of ECoG data obtained from five participants that took part in a battery of speech production tasks: Auditory Repetition (AR), Auditory Naming (AN), Sentence Completion (SC), Word Reading (WR) and Picture Naming (PN).These were designed to elicit the same set of spoken words across tasks while varying the stimulus modality [41] and provided 50 repeated unique words (400-800 total trials per participant) all of which were analyzed locked to the onset of speech production. We start with an overview of our speech decoding approach.

### 2.1 Speech Decoding Approach

- **ECoG Decoder**. The decoder maps the ECoG signals to a set of speech parameters (describing both the voiced and unvoiced components) which are then synthesized to speech spectrograms (Figure 1). The ECoG decoder architecture is based on recent advances in convolutional neural networks leveraging the ResNet approach [24]. We construct a modified ResNet model with nine layers that treat the cortical input as a spatiotemporal three-dimensional tensor (two dimensions for the electrode array and one for time, see Methods for details). The decoder is trained such that its output parameters match the reference parameters derived from a speech encoder (which is learnt separately in an unsupervised manner). Furthermore, our approach ensures that the speech spectrogram derived from these parameters and constructed by the speech synthesizer matches with the actual speech spectrogram. We use this approach to be more data-efficient and allow us to train on a small set of samples for each patient.
- **Speech Parameters**. Our speech representation is motivated by the vocoders used for low-bit-rate speech compression dating back to the 1980s. We model speech signals as a mixture of voiced and unvoiced components, with the voiced component described by a source-filter model (dynamically filtered harmonic signals) [13] and the unvoiced component generated by white noise broadband filtering. In addition to the mixing parameter, our representation includes speech formant information (frequency, bandwidth, etc.) and loudness (i.e., the energy of speech). See Methods Figure 6 for details.
- **Synthesizer**. We use a set of signal processing equations (such as harmonic oscillation, noise generation, filtering, etc.) to synthesize the spectrogram from our proposed speech parameters. We can train the ECoG decoder with a limited amount of training data by limiting the number of speech parameters and using differentiable signal processing equations. It is noteworthy that the equations we use are differentiable (see Differentiable Speech Synthesizer in Extended Data A.1), which allows for backpropagation from the spectrogram to the actual learning of the decoder.
- **Speech Encoder**. The speech encoder is pre-trained using an independent unsupervised approach before the ECoG decoder training. The encoder is trained to generate a set of speech parameters from a given spectrogram, from which the aforementioned speech synthesizer can reproduce the spectrogram. This pre-trained encoder generates reference speech parameters from actual speech signals used for the training of the ECoG decoder. The unsupervised process can be easily used to train the speech encoder from any set of speech signals, including patient-specific speech (see details in the Method section 4.4 and Extended Data A.1). Importantly, this process constrains the speech parameter space to optimize the training of our ECoG decoder, and the parameters can directly drive a speech synthesizer based on differential equations.

We trained three separate models using the proposed pipeline, varying in the causality of the temporal convolution used in the ECoG decoder. The causal model uses only past (up to the current) neural signals to produce the current speech sample, which reflects feedforward processes. The anticausal model only uses current and future neural signals, reflecting feedback processes. And finally, the non-causal model uses both the past, current, and future neural signals, which are typically used in previous literature and confounds feedforward and feedback processing. The causal and anticausal models allow us to directly assess and tease apart the feedforward and feedback contributions of different cortical regions. It is important to recognize that only causal models are appropriate for real-time speech prosthetic applications.

### 2.2 Speech decoding performance

We first demonstrate that our approach produces accurate speech decoding with detailed acoustic features. The model’s decoded spectrogram preserves the spectro-temporal structure of the original speech. It reconstructs both vowels, consonants (Figure 2a) as well as the overall spectral energy distribution (Extended Data Figure E1). These acoustic details result in a reconstruction that preserves the speakers’ timbre (see Supplementary Video) and leads to naturalistic voice decoding. Our model’s speech parameters which include loudness, formant frequency, and the mixing parameter (i.e., the relative weighting between voiced and unvoiced components), are decoded accurately with the correct temporal alignment of each word onset and offset (Figure 2b, c). The overall accuracy of the fundamental frequency (i.e., pitch), the first two modeled formants (i.e., F1, F2), and the transition between voiced and unvoiced sounds are a major driving force for accurate speech decoding as well as naturalistic reconstruction that mimics the patient’s voice.

**Figure 2:**
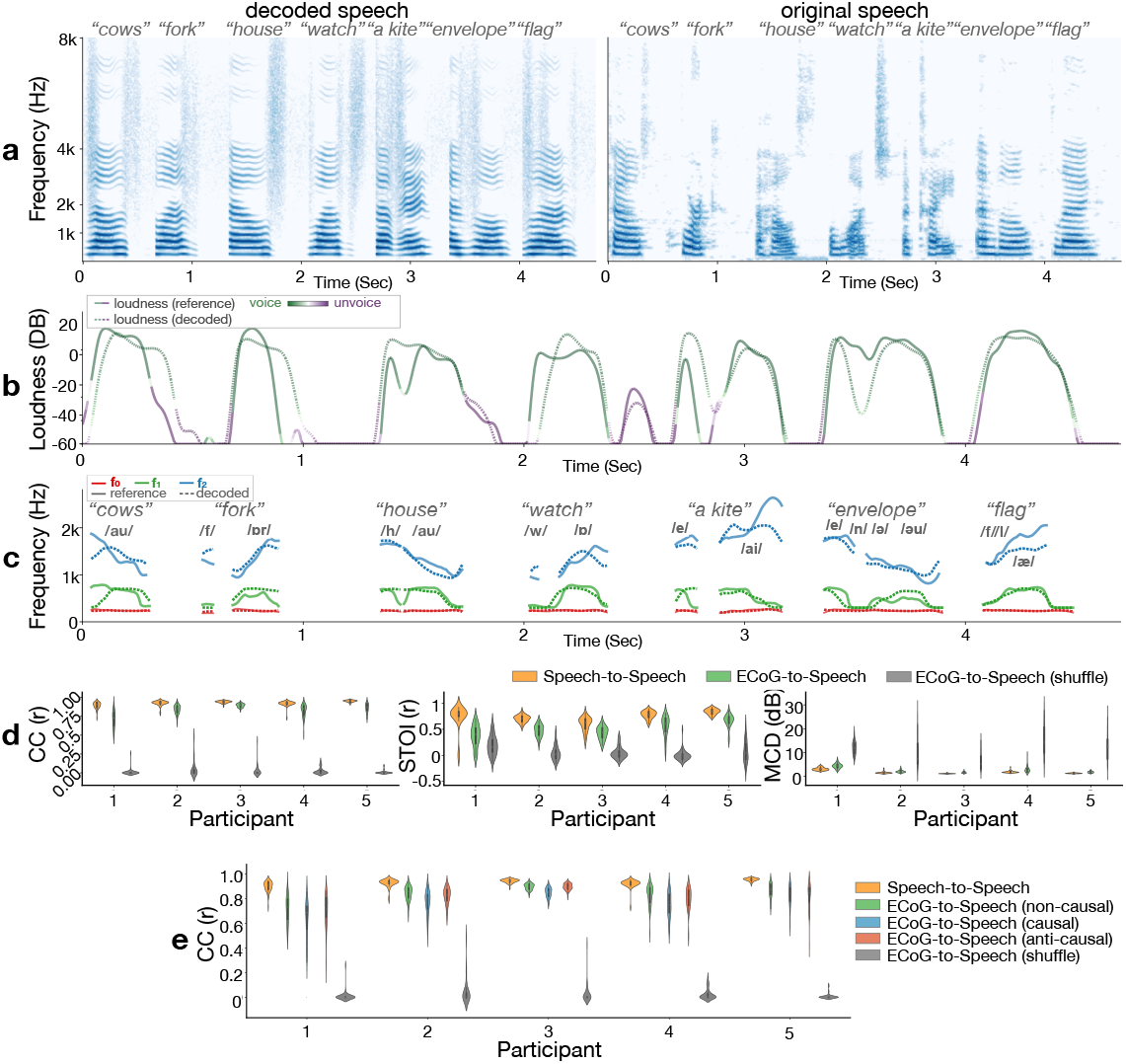
Comparison of original and decoded speech produced by the model. (a) Spectrograms of decoded (left) and original (right) speech exemplar words. (b) Decoded loudness parameter with the voiced (mostly vowel) or unvoiced (mostly consonant) mixing parameter color-coded over the loudness curves. The same color spread and amplitude trend between decoded (dashed) and reference (solid) curves reflect accurate decoding of voice and unvoiced phonemes with correct energy and temporal alignment. (c) Frequencies of the first two formants (F1, F2) and the pitch (F0). The matching between decoded (dashed) curves and reference (solid) curves in both frequencies during each phoneme and the overall temporal dynamic leads to intelligible and naturalistic decoding of voiced sounds. (d) Evaluation of the decoded speech quality in objective metrics. The correlation coefficient of spectrograms (CC, left), short-time objective intelligibility (STOI, middle), and Mel cepstral distortion (MCD, right) are used for the evaluation. Note that lower MCD values represent better performance. Both the reconstructed speech from the speech auto-encoder (yellow) and the speech decoded by the ECoG decoder (green) are reported. Additionally, the performance of a model trained on shuffled data (trained by matching the decoded spectrogram from the neural signal in a given duration to a randomly selected segment of spectrograms during the entire recording session) is also reported as a control. (e) Comparison of the CC metric among noncausal (green), causal (blue), and anticausal (red) models. Compared to the shuffled model (the same shuffled model as in Figure 2d), the comparable performance across noncausal, causal, and anticausal models demonstrates sufficient information for decoding speech from both feedforward and feedback signals during speech production.

In order to evaluate the performance and quality of speech, we used several objective metrics, including the correlation coefficient (CC) between the decoded spectrogram and actual produced speech [2, 3, 25], an objective measure for speech intelligibility known as the Short-Time Objective Intelligibility (STOI) [3, 45], and a measure of spectral distortion, Mel-cepstral distortion (MCD) [4, 35]. Across all participants and metrics, our neural decoding results performed well above chance (Figure 2d in grey; estimated using shuffled data, see Methods section 4.6) and approached an upper bound of performance based on the unsupervised autoencoder (i.e., speech-to-speech) which did not use neural data. The performance range across metrics, and our participants were equal to and often better than the current literature [2–4, 25]. Critically, all these models represent the non-causal case (Figure 2d) that uses data both from the past (feedforward) and the future (feedback), as is currently a common practice [1–4, 37] except a nominal few models [25].

In order to directly assess the performance of the causal (predicting using only the past) and anticausal (predicting using the future feedback) models and compare them with the non-causal (using past and future) model, which is standard in the field, we trained three separate models varying the temporal convolution direction. Our results (Figure 2e) show a slight decrease in performance with the causal model. However, it performs close to the other models while providing a causal interpretation, which only uses past signals to predict future speech. This is encouraging, as it suggests that, with additional improvement in the decoder design and training, it is possible to design practically applicable neuroprosthetic speech synthesizers. Also, comparable performance between causal, anticausal and non-causal approaches indicates a similar amount of information contained by feedforward and feedback signals. Both causal and anticausal models are appropriate for feedforward-feedback analysis and comparison.

### 2.3 Feedforward and feedback cortical contributions to speech production

To elucidate the feedforward and feedback contribution of different cortical regions to speech production, we examined the relative contribution of each electrode to decoding speech in our models. We derived the relative contribution by quantifying how the input signal at a particular electrode affects the overall accuracy (measured by the CC) of the reconstructed speech in the causal and anticausal models, respectively (see Methods 4.5). In both the causal and anticausal models, peri-sylvian electrodes were important for speech decoding; however, there was a surprising recruitment of frontal regions when decoding speech based on the feedback (anticausal model, Figure 3b) as well as recruitment of temporal sites when decoding speech based on the feedforward signals (causal model, Figure 3c). We only show significant contributions that are above a threshold derived from the shuffled model (depicted in Figure 3d). In order to quantify the prevalence of feedforward or feedback processing, we directly contrasted the two and projected the results onto the cortex (Figure 3e). To ascertain regions that contribute significantly more to feedback or feedforward processing, we conducted a region of interest analysis, based on within-subject anatomical labels of each electrode (see Methods section 4.3), testing for an increase in causal or anticausal contributions across trials (non-parametric paired Wilcoxon test; Figure 3f). We found a surprisingly mixed distribution of causal and anticausal contributions within both temporal and frontal cortices. A majority of temporal cortex were predominantly anticausal, including caudal superior temporal gyrus (STG; Wilcoxon sign rank, P=1.607E-15, Z=9.6234) and portions of middle temporal gyrus (MTG; rostral MTG: Wilcoxon sign rank test P=2.5108E-04, Z=4.9359, and middle MTG: Wilcoxon sign rank test P=1.5257E-13, Z=9.0185) as well as supramarginal cortex (Wilcoxon sign rank test P=1.1144E-04, Z=5.3919), implicating it in processing the auditory feed-back signals for speech production. However, there was also a significant causal contribution in rostral STG (Wilcoxon sign rank test P=0.0332, Z=-2.9628). Similarly, the majority of sensorimotor cortex was predominantly casual, implicating it in processing the motor speech commands including ventral precentral (Wilcoxon sign rank, P=4.9511E-08, Z=-7.1409) and postcentral gyri (Wilcoxon sign rank, P=6.419E-04, Z=-4.9612). However, the dorsal division of precentral gyrus was equally causal and anticausal (Wilcoxon sign rank, P=0.4349, Z=0.6525), implicating it in processing both feedforward and feedback information equally. Within the inferior frontal cortex, we found a striking division of function wherein pars opercularis was significantly causal (Wilcoxon sign rank test, P=8.0693E-15, Z=-9.6185) while pars triangularis was significantly anticausal (Wilcoxon sign rank test, P=2.6715E-06, Z=6.3518). Overall, these findings provide evidence for a mixed feedforward and feedback processing of speech commands and their reafference across temporal and frontal cortices, in contrast to a dichotomous view.

**Figure 3:**
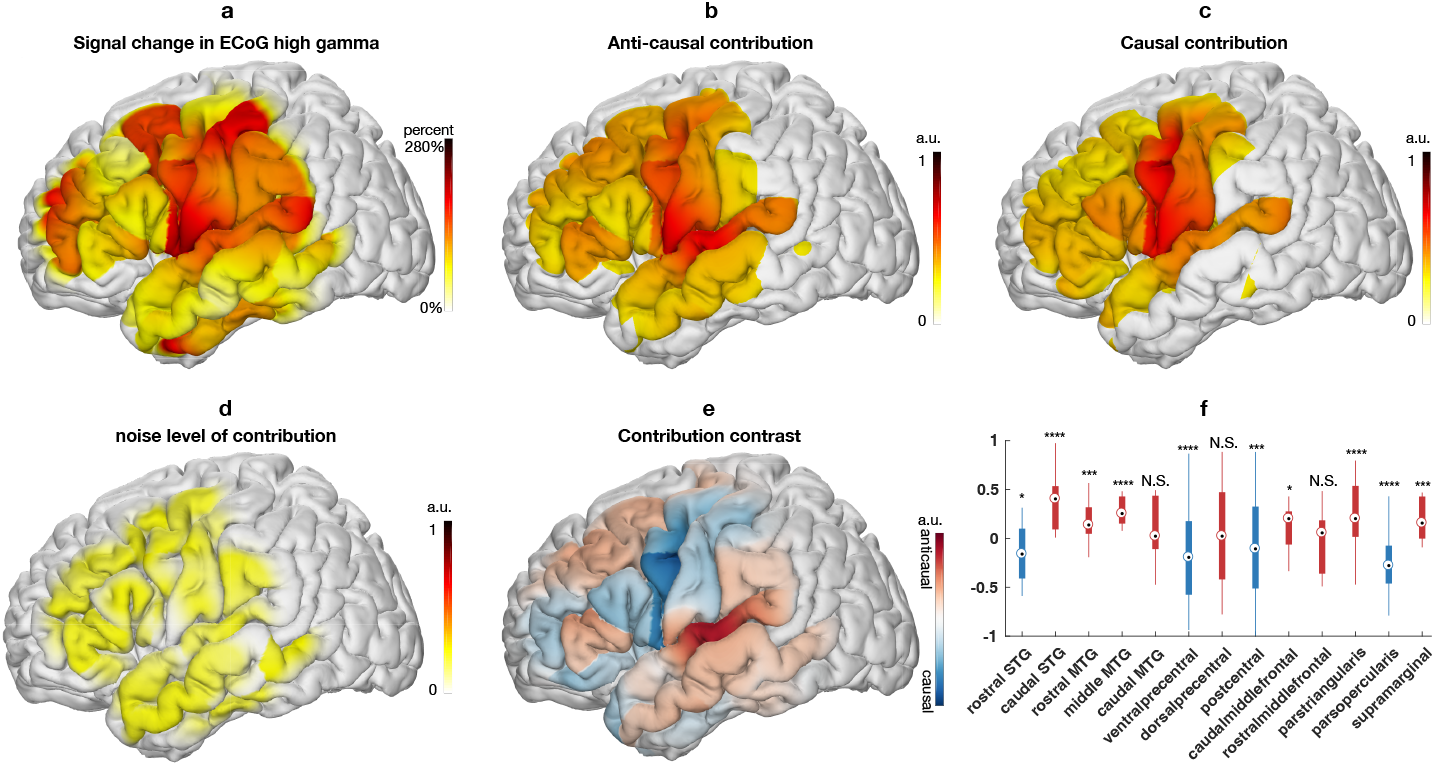
(a) averaged signal of input ECoG projected on the standardized MNI anatomical map. The colors reflect the percentage change of high gamma compared to the baseline level during the pre-stimulus baseline period. (b) shows the anticausal contribution of different cortical locations (red indicates higher contribution), while (c) illustrates the causal contribution. (d) noise level of the contribution analysis evaluated by the contributions from the shuffled model. Contributions below noise level are not shown in (b) and (c). (e) the contrast obtained by taking the difference of the anticausal and causal contribution maps (red means higher anticausal contribution, while blue means higher causal contribution). The boxplots (f) show the average difference in each cortical region (*: P-value<0.05, **: P-value<0.01,***: P-value<0.001,****: P-value<0.0001).

### 2.4 Temporal dynamics and receptive fields of speech production

Speech production includes articulatory planning and executing the motor commands, processes that recruit distinct regions of frontal cortex [16]. However, their exact temporal receptive fields remain poorly understood. Earlier, we examined the causal and anticausal cortical contributions during speech articulation. Next, we examine articulatory planning and articulation of speech production stages and derive the related temporal receptive fields across the cortex. We leverage the receptive fields to test how cortical regions contribute differently to speech decoding with time and how frontal cortex dynamics change when feedback is introduced (after articulation starts). Both feedforward and feedback information is processed in tandem.

We employed a similar occlusion approach to derive the temporal receptive fields as in the previous section. However, we quantified how the input signal at a particular electrode affects the accuracy of the reconstructed speech across varying delays (see Methods section 4.7). This approach allowed us to quantify the contribution of a specific electrode in the model as a function of delay relative to speech decoding, similarly to classical temporal receptive fields (i.e., TRF). We conducted this analysis for both causal and anticausal models during two epochs – one prior to production (−512ms ~ 0 ms; Figure 4a) and the other during production, which included both causal and anticausal components (0ms ~ 512ms; Figure 4b, c). The projection of all the temporal receptive fields onto the cortex, which were significantly above a threshold derived from the shuffled model, are plotted in Figure 4 as a function of delay. We found an increased frontal and MTG contribution prior to production (Figure 4a) compared with during production (Figure 4b). These processes are likely related to articulatory planning and lexical retrieval prior to speech production. During production, there was a prominent sharpening of ventral precentral gyrus receptive fields marked by a significant increase in contribution compared with pre-production (Wilcoxon sign rank test, P=8.3979E-05, Z=5.4203). While a majority of prefrontal regions engaged prior to production, there was a significant decrease in contribution across pars triangularis (Wilcoxon sign rank test, P=1.8493E-32, Z=-13.6074), middle frontal gyri (MFG; Wilcoxon sign rank test, P=3.9177E-09, Z=-7.6103 for caudal and P=4.1581E-04, Z=-4.8311 for rostral) except for pars opercularis (Wilcoxon sing rank test, P=0.4819, Z=0.2066). Similarly, to our previous results (Figure 3e,f), during production, we found a significant increase in anticausal contribution for caudal STG (Wilcoxon sign rank test, P=2.6789E-17, Z=9.6711), pars triangularis (Wilcoxon sign rank test, P=0.0162, Z=3.9003) and caudal MFG (Wilcoxon sign rank test, P=0.0045, Z=3.9862) compared with causal contributions. This confirms the anatomical-functional division of the inferior and middle frontal gyri as well as caudal (Wilcoxon sign rank test, P = 2.6789E-17, Z = 9.6711) and rostral separation of STG (Wilcoxon sign rank test, p= 0.0343, Z= −2.9457).

**Figure 4:**
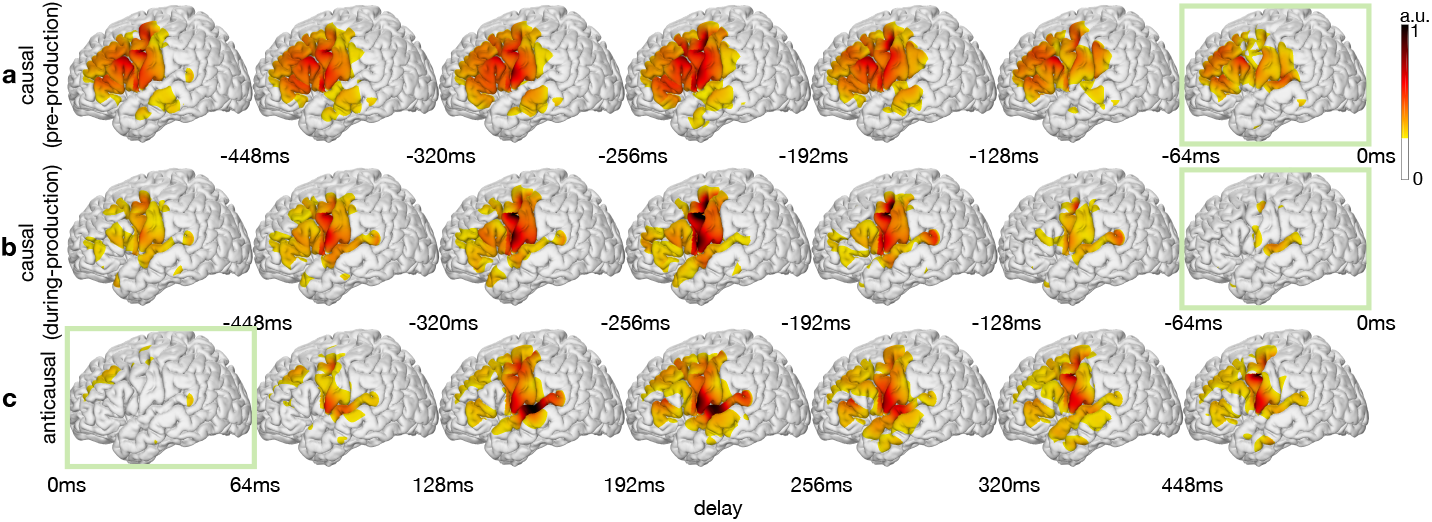
Spatial-temporal receptive fields based on decoding contribution. The contribution to decoding the current speech from cortical neural responses with certain temporal delays. (a) and (b) are the feedforward spatial-temporal receptive fields derived from the causal model by evaluating the contribution of past (negative delays) neural signals during a period before production onset (a) and after onset (b). (c) represents the feedback spatial-temporal receptive fields derived from the anticausal models that evaluate the contribution of future (positive delays) neural signals during feedback after articulation. Contributions below significance (pꜟ0.05) representing the noise level are clipped and not shown in the plots.

Next, we conducted a region of interest analysis, based on within-subject anatomical labels of each electrode, in order to derive the temporal receptive curves per region (Figure 5). This approach provides critical insight as to the temporal tuning and peak recruitment of various regions to feedforward processing prior to (Figure 5a) and during production (Figure 5b) as well as feedback processing (Figure 5c). We found a shift in receptive field tuning for the two subdivisions of precentral gyrus. Prior to production, dorsal and ventral precentral gyri were not significantly different from each other (Wilcoxon sign rank test, P=0.454, Z=-0.36103), and had close peak times (−196ms, −192ms prior to speech for ventral and dorsal precentral gyri, respectively). However, during production, these dynamics shifted and we found a significant decrease in dorsal precentral causal contribution (Wilcoxon sign rank test, P=4.7575E-05, Z=-5.6272) accompanied by a temporal separation of peaks (−208ms, −184ms for ventral and dorsal precentral gyri, respectively; Figure 5a,b). Within the inferior frontal gyrus, we found pars opercularis was recruited similarly both prior to production and during production for feedforward processing (Wilcoxon sign rank test, P=0.5922, Z=1.7462) at a peak delay of −248ms and −280ms, respectively. During production, pars triangularis had a selective increase in recruitment for anticausal compared with causal contributions (Wilcoxon sign rank test, P=0.0162, Z=3.9003), implicating it in increased feedback processing (Figure 4c, Extended Data Tables 2, 3). The anticausal receptive fields during production provide evidence for feedback processing most strongly contributed by caudal STG, with the earliest peak in contributions seen in dorsal precentral gyrus (144 ms) and caudal STG (168 ms) followed by parietal (supramarginal 184ms, postcentral 192ms) and ventral precentral (280 ms) gyri ( Extended Data Table 3). These findings suggest a preferential recruitment of prefrontal cortices in feedforward processing prior to production followed by a shift in dynamics during production when feedforward and feedback signals are jointly processed with anatomical divisions of labor.

**Figure 5:**
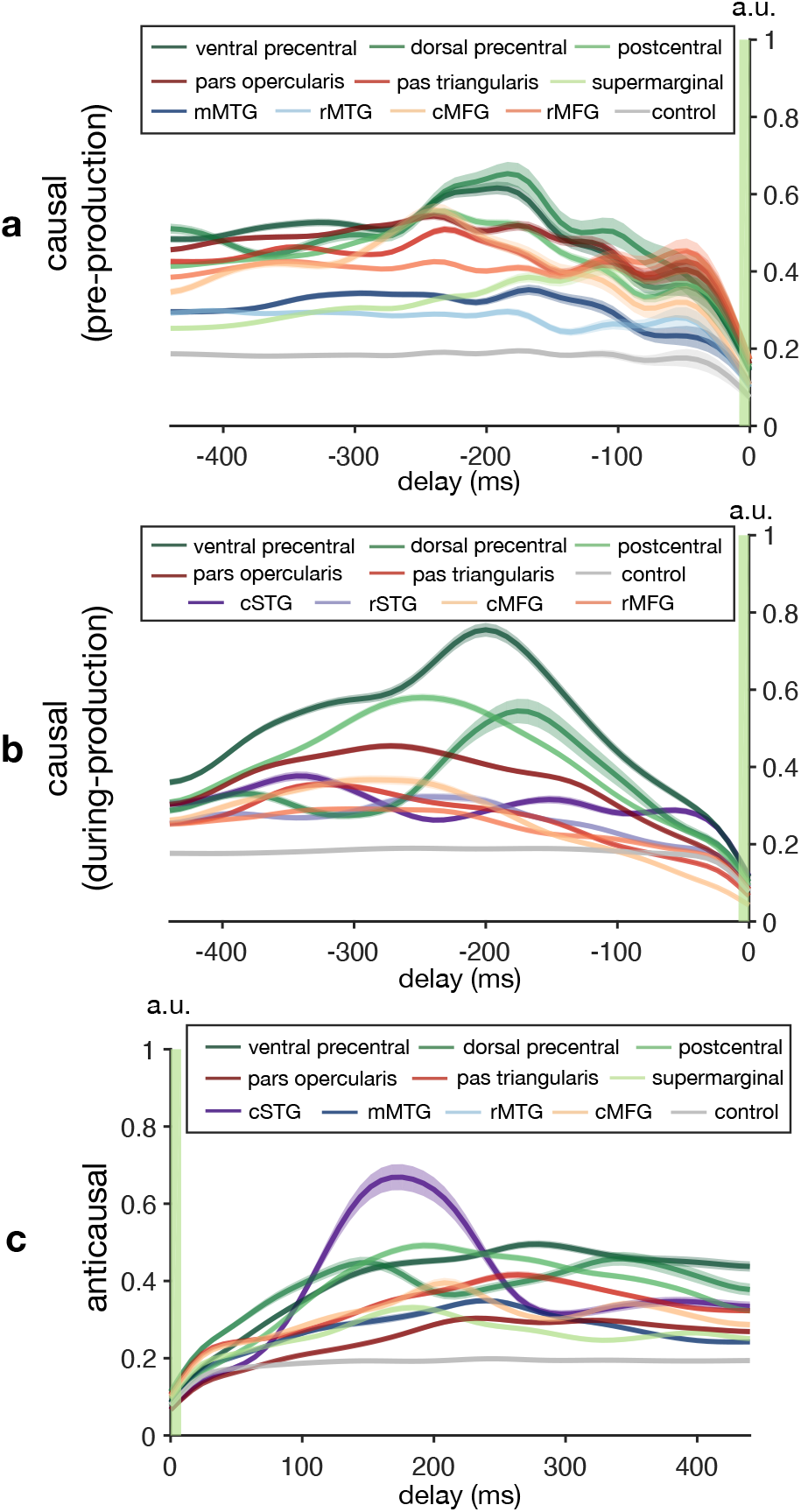
The temporal receptive field across anatomical regions. The contribution to decoding the current speech from cortical neural responses with certain temporal delays. (a) and (b) are the feedforward temporal receptive fields derived from the causal model by evaluating the contribution of past (negative delays) neural signals during a period before production onset (a) and after onset (b). (c) represents the feedback temporal receptive fields derived from the anticausal models that evaluate the contribution of future (positive delays) neural signals during feedback after articulation. The temporal propagation of the shuffled model estimates the noise level dynamics (grey curves in plots). Only regions significantly above noise level (Wilcoxon sign rank test on across-time averaged data, P<0.05) are reported.

**Figure 6:**
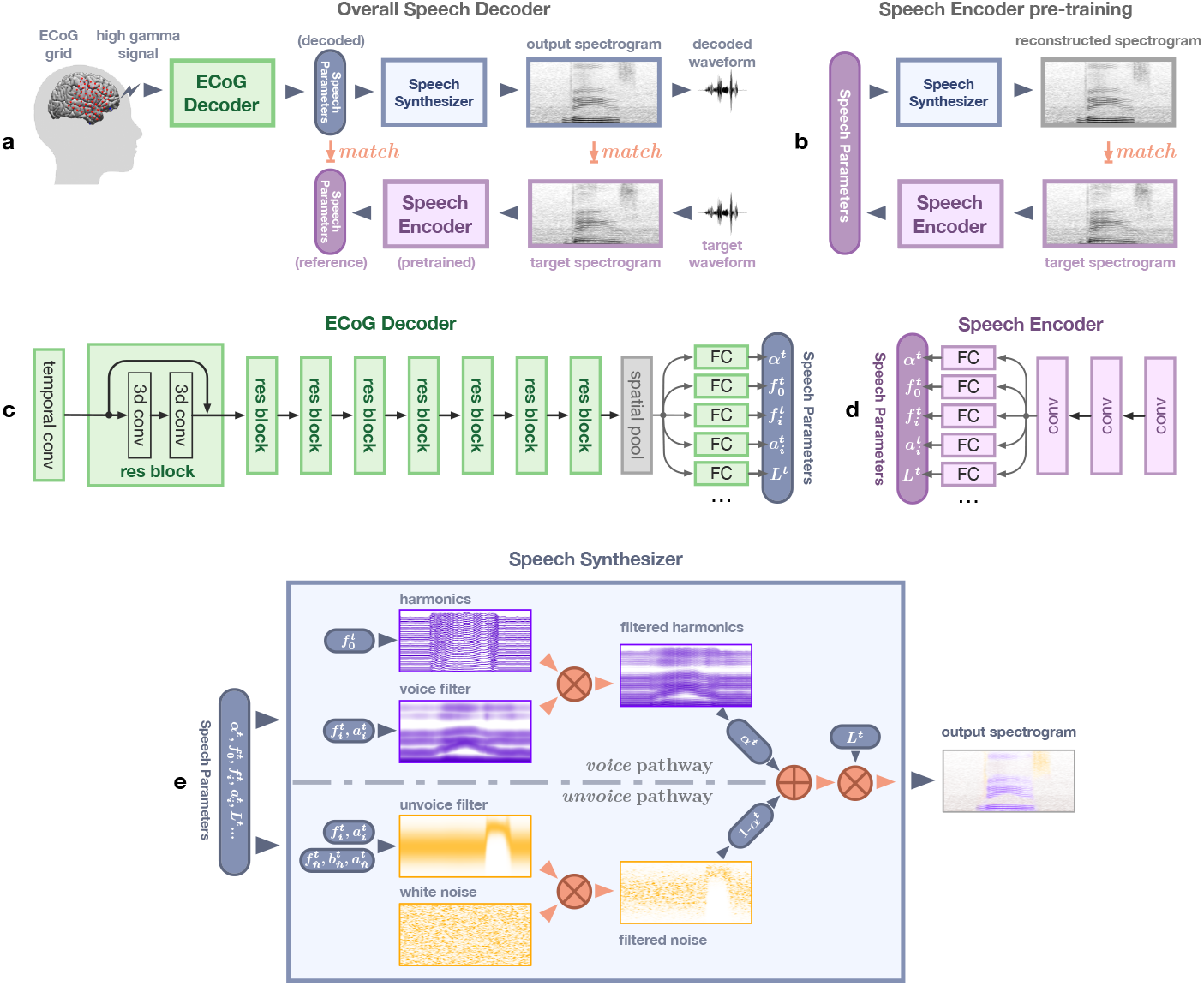
Structure of the decoding framework. (a) The overall network architecture (same as in Figure 1, repeated here for ease of understanding of the architecture). (b) The auto-encoder used to pretrain the speech encoder. The speech encoder is trained to generate proper speech parameters that can reconstruct input spectrograms through the speech synthesizer. (c) The ECoG decoder is a modified spatio-temporal residual network. After an initial temporal convolutional layer and eight residual blocks (constructed by three-dimensional convolution layers), multiple subnetworks (using one or two fully connected layers) generate speech parameters separately. (d) The speech encoder in (b) has three convolutional layers followed by the same multi-head output structure as in (c). (e) Illustrates the processes within the speech synthesizer. The harmonics (in voice pathway) and white noise (unvoice pathway) are generated and filtered (multiplication in spectrogram domain) by voice and unvoice filters, respectively. The filtered results are then weighted averaged according to the mixing parameter and then amplified by the loudness parameter.

## 3 DISCUSSION

Our study leverages a novel deep learning approach together with neurosurgical recordings and, to our knowledge, is the first to dissociate direct feedforward and feedback cortical contributions during speech production. Our neural network architecture achieves state-of-the-art decoding of speech production, by tapping an interpretable compact speech representation and can be altered to focus on causal, anticausal and non-causal decoding. Our analyses of the cortical contributions driving the performance of these models reveal a mixed distribution of feedforward and feedback processing during speech production. This was prominent in inferior, middle frontal, and superior temporal gyri which exhibited an anatomical division between feedforward and feedback processing. Lastly, we show a change in the temporal dynamics of frontal recruitment during speech planning through production, characterized by a shift of inferior frontal and precentral gyri recruitment, processing both feedforward and feedback information at different time points and spatial locations.

A growing number of studies have leveraged deep neural networks for cortical speech decoding. Convolutional neural networks (CNN) [1, 3, 46, 47] and recurrent neural networks (RNN) [4] have mapped ECoG signals into speech and text [37]. However, our approach diverges from these studies. *Firstly*, we develop a novel differentiable speech synthesizer that can generate natural speech from a compact set of interpretable speech parameters based on several signal processing equations. This rule-based synthesizer allows for unsupervised pre-training of meaningful encoded representations (reference speech parameters), as well as reduces the capacity of the entire model and increases training data efficiency. Our approach provides a direct mapping to a patient’s voice. It eliminates the need for labeled articulatory data that maps speech to articulatory dynamics as proposed by Anumanchipalli et al. [4]. *Secondly*, our compact speech representation leverages an interpretable decomposition of speech into voiced and unvoiced components. This decomposition is biologically necessary, has been reported in neural representations across frontal and temporal cortices [8, 31] and stands in contrast to other traditional speech synthesizing approaches [13, 14]. *Lastly*, the speech neural decoding models to date mostly employ non-causal operations. Since such decoders require both past and future information for decoding, they are not applicable for real-time speech prosthetic application. Further, mixed operations hinder disentangling feedforward and feedback cortical contributions. In addition to providing a causal model which directly translates to practical speech prosthetics, our approach provides one of the first reports that can dissociate feedforward and feedback cortical contributions during speech production.

During speech production, we process feedforward and feedback signals in tandem. It was previously impossible to disentangle the two. Attempts have focused on experimental manipulations which change the feedback by shifting frequency [6] or time [39]. However, these manipulations change the cortical dynamics and introduce other cognitive processes due to hearing one’s own voice altered as well as induced motor compensation. We applied convolution filters with different causality to directly train models to disentangle feedforward (i.e., causal models) and feedback (i.e., anticausal models) contributions of cortical regions. Feedforward and feedback processes are critical for driving articulatory vocal tract movement. The feedforward pathway generates an initial articulatory command and predicts sensory (auditory and somatosensory) targets; the feedback pathway compares the targets with the perceived sensory feedback and updates subsequent feedforward commands. The exact mapping between anatomical regions and their contribution to specific functional roles differ across speech motor control models ( [23], [30]). Further, these findings have been developed based mostly on non-invasive studies which have low temporal (e.g., fMRI) or spatial resolution (e.g., M/EEG). Our high spatio-temporal resolution ECoG data together with advanced deep neural networks provides a fine-grained mapping of the cortical feedforward and feedback speech networks.

Consistent with the predominant speech motor control models, our results showed a dominant feedforward process in the ventral motor and pars opercularis of the inferior frontal gyrus, while posterior superior temporal and supramarginal gyri in the parietal lobe showed feedback. However, in contrast to these models, we found that cortices in the frontal lobe, including pars triangularis and caudal middle frontal, are predominantly feedback in nature, while rostral STG appears feedforward. This feedback processing across frontal cortices became even stronger when we limited our analyses to the speech production epoch (Figure 4c, Extended Data Table 3). Additionally, most gyri (inferior frontal, caudal middle frontal, superior temporal, precentral, and postcentral cortices, see Extended Data Table 2) had both feedforward and feedback contributions above the noise level derived from the shuffled model, suggesting the feedforward and feedback processing can mix in these regions.

Our results highlight the anticausal feedback signature exhibited by sensorimotor and frontal cortices. While this goes against the canonical model of the frontal cortex in an action-perception loop [18], our findings complement a growing body of evidence showing specific responses in the frontal cortex to auditory stimuli during perception. Cheung et al. [9] found distinct auditory receptive fields as well as robust passive listening responses in ventral precentral gyrus. Similarly, the dorsal division of precentral gyrus has recently been implicated in processing auditory feedback of altered speech as well as responding robustly during passive listening [39]. However, this begs the question as to why the speech motor cortex is processing auditory information. Our feedback contribution analysis suggests that the auditory processing is specifically leveraged for anticausal processing of the reafferent signals during production. Indeed, our results show that dorsal precentral gyrus decreases feedforward processing while engaged in actual speech production (Figure 5b) and is recruited for feedback at an early time point together with temporal cortices (Figure 5c). Under this view, the auditory frontal responses seen during passive listening may constitute a representation dedicated to feedback processing when speech is produced.

To summarize, we provided a new approach to decode speech production and interrogate the recalcitrant problem of mixed feedforward and feedback processing during speech production. We were able to leverage feedforward processing only in causal models to drive neural speech prosthetics (as opposed to the literature using non-causal processing [1–4, 37]) as well as provide insights into the underpinning cortical drivers. Our results suggest a mixed cortical architecture in frontal and temporal cortices that dynamically shifts and processes both feedforward and feedback signals across the cortex in contrast to previous views associating feedforward or feedback processing of speech with primarily anterior and posterior cortices, respectively.

## 4 METHODS

### 4.1 Participants and experiments

The brain activity data were obtained from five patients, including three female and two male native English speakers, undergoing treatment for refractory Epilepsy at NYU Langone hospital, with implanted electrocorticographic (ECoG) subdural electrode grids. All experimental procedures were approved by the NYU Grossman School of Medicine Institutional Review Board. Patients were provided written and oral consent at least one week prior to surgery by a research team member after separate consultation with the clinical care provider. The subjects were instructed to complete five tasks to pronounce the target words in response to certain auditory or visual stimuli. The five tasks were:

- Auditory Repetition (AR, i.e., to repeat the auditory words).
- Auditory Naming (AN, i.e., name a word based on an auditory presented definition sentence).
- Sentence Completion (SC, i.e., complete the last word of an auditorily presented sentence).
- Visual Reading (VR, i.e., read aloud visually presented word in written form).
- Picture Naming (PN, i.e., naming a word based on a visually presented color line drawing).

Each task contained the same 50 unique target words while varying stimulus modalities (auditory, visual, etc.). Each word appeared once in the AN and SC tasks and twice in the other tasks. For Participants 1-3, the five tasks included 400 trials of the produced words and the corresponding ECoG recordings. The produced speech in each trial has an average duration of 500 ms. We repeated the same five tasks twice for Participant 4 and collected data from 800 trials. For Participant 5, we collected 800 trials by repeating the tasks twice, and we also ran an additional AR task (200 trials) which provided 1000 trials in total.

### 4.2 Data collection and preprocessing

A microphone recorded the subject’s speech during the tasks and was synchronized to the clinical Neuroworks Quantum Amplifier (Natus Biomedical, Appleton, WI), which records ECoG. The recordings sampled peri-sylvian cortex, including STG, IFG, pre-central, and postcentral gyri. The ECoG implanted array included standard 64 clinical 8×8 macro contacts (10 mm spacing) as well as 64 additional interspersed smaller electrodes (1 mm) between the macro contacts (providing 10 mm center-to-center spacing between macro contacts and 5 mm center-to-center spacing between micro/macro contacts, PMT corporation, Chanassen, MN). This FDA-approved array was manufactured for the study, and a member of the research team explained to patients that the additional contacts were for research purposes during consent. The ECoG arrays were implanted on the left hemisphere in all participants’ brains and placement location was solely dictated by clinical care. We trained separate sets of decoding models for each participant. We randomly selected 50 out of all trials from the five tasks for testing and used the remaining data for training. The results reported are for testing data.

Each electrode sampled ECoG signals at 2048 Hz, which was decimated to 512 Hz prior to processing. After rejecting electrodes with artifacts (i.e., line noise, poor contact with cortex, and high amplitude shifts), we subtracted a common average reference (across all valid electrodes and time) from each individual electrode. Electrodes with inter-ictal and epileptiform activity were removed from the analysis (note that the large number of temporal electrodes were removed from patients 4 and 5 for this reason). We then extracted the envelope of the high gamma (70-150 Hz) component from the raw signal with the Hilbert transform and further downsampled it to 125 Hz. The signal of each electrode over the silent baseline of 250 ms before the stimulus was used as the reference signal, and each electrode’s signal was normalized to the reference mean and variance (i.e., z-score).

### 4.3 Electrode localization

Electrode localization in subject space, as well as MNI space, was based on coregistering a preoperative (no electrodes) and postoperative (with electrodes) structural MRI (in some cases, a postoperative CT was employed depending on clinical requirements) using a rigid-body transformation. Electrodes were then projected to the surface of the cortex (preoperative segmented surface) to correct for edema-induced shifts following previous procedures [48] (registration to MNI space was based on a non-linear DARTEL algorithm). Based on the subject’s preoperative MRI, the automated FreeSurfer segmentation (Destrieux) is used for labeling within subject anatomical locations of electrodes.

### 4.4 Speech decoding framework

The backbone of our neural decoding framework is constructed by an ECoG decoder and a speech synthesizer (Figure 6a or Figure 1). During testing, from the high gamma components of the ECoG signal, the decoder generates a set of speech parameters that drive a differentiable speech synthesizer to generate speech spectrograms (and corresponding waveforms by the Griffin-Lim algorithm). Besides being trained to work with the speech synthesizer to output spectrograms matching the target spectrograms, the ECoG decoder is also trained to match its output with a set of reference speech parameters. This reference matching training strategy provides a more direct gradient to the ECoG decoder such that it converges faster and is less prone to overfitting.

The reference speech parameters are derived from a pre-trained speech encoder. During pre-training, the speech encoder and the speech synthesizer fulfill an auto-encoding task (i.e., mapping the input spectrogram to the speech parameters and back to the spectrogram) (Figure 6b). When such speech-to-speech reconstruction is accurate, the parameters generated by the speech encoder should provide physically meaningful speech parameters. Since the pre-training is unsupervised and the subject speech audio data is easy to collect, obtaining the reference speech parameters is straightforward. Note that the speech-to-speech autoencoder and the reference parameters are only used for the training of the ECoG decoder. Once the ECoG decoder is trained, the trained decoder and the speech synthesizer can be used to convert ECoG signals to speech without the need for reference parameters.

More details of the structure of the speech synthesizer (Figure 6e), ECoG decoder (Figure 6c), Speech encoder (Figure 6d), and loss can be found in Extended Data A.1.3.

### 4.5 Revealing delay-dependent contribution of different cortical regions from the trained ECoG to speech model

Before formally defining the various contribution scores, we introduce the following notations: A_ref_[*s*]: the reference spectrogram over a time duration S centered at time *s*, i.e., from *s* – S/2 to *s* + S/2, derived by the speech-to-speech autoencoder. A_intact_[*s*]: the model output with “intact” input (i.e, all ECoG signals are used). 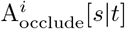: the model output at time duration centered at s when the *i*th ECoG electrode signal in the time duration centered at t from 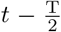 to 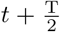 is occluded. *r*(·, ·): correlation coefficient between two signals. We define the contribution of *i*th electrode in time duration centered at *t* to the output over duration centered at *s* by the reduction in the correlation coefficient between the output signal with the reference signal over the duration *s* when the ith electrode signal in duration *t* is occluded. Specifically:

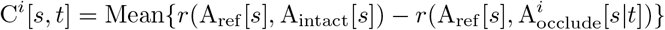

where Mean{·} denotes averaging across all testing samples.

To generate the contribution map, we first determine the contribution of each electrode (with a corresponding location in the MNI coordinate), which is then diffused into the surrounding area in the same anatomical region using a Gaussian kernel. Since our ECoG grid has hybrid density, to remove the effect of non-uniform density on the diffused result, we normalize the result of each region by the local grid density. The results shown in Figures 3,4, and 5 are obtained by averaging the contribution maps obtained for all test samples for all participants.

### 4.6 Visualizing spatial contribution map

The contribution of the entire signal at the *i*-th electrode to the entire output signal, C^*i*^, is obtained by using the method in Section 4.5 with S and T covering the entire input and output signal duration. The causal and anticausal contribution plots in Figure 3 are generated by applying such analysis to the learned anticausal model (Figure 3b) and causal model (Figure 3c), respectively. The contrast of the anticausal and causal contribution (Figure 3e) for each is the difference between the causal and anticausal contribution map. The noise level for the contribution analysis (Figure 3d) is generated from the shuffled model using non-causal processing (the shuffled model is trained on an artificial dataset with temporal misaligned input-output, and hence models of different causality are equivalent). To generate per region feedback-feedforward box plot (Figure 3f), we calculate the contrast contributions averaged over electrodes of the same within-subject anatomical labels corresponding to each region.

The contrast of the anticausal and causal contribution (as is shown in Figure 3e) of electrode *i* is defined as

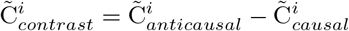

In order to examine electrode polarization to anticausal or causal contribution, we calculate the normalized version of anticausal and causal contribution contrast:

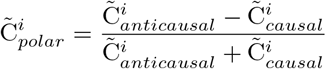

By normalizing the contrast of anticausal and causal contribution, 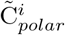 emphasize the angle of contribution directing towards anticausal or causal, rather than their contrast. This is what is visualized in Figure E2 a,b in Extended Data (only for those electrodes with either anticausal contribution attribute 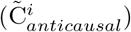 or causal contribution attribute 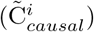 above noise level determined by the shuffled model). This is what is shown in Extended Data Figure E2.

### 4.7 Visualizing spatial-temporal contribution receptive field

When evaluating the contribution over a finite duration we use small temporal scope S = T = 64ms. To Evaluate the contribution of an electrode signal to the output with various delay, denoted by *τ*, we average C^*i*^[*s*, *s* + *τ*] for all s in a certain duration leading to

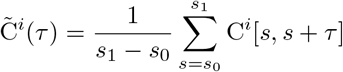

Here we assume the effect of delay is independent of actual output time *s*. When *τ* ≤ 0, 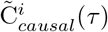 reveals the causal contribution of electrode *i* to the output (Figure 4 a,b). To investigate pre-production contribution, we restrict *s* + *τ* and *s* to be no later than the onset of production (vise-versa for during-production analysis). When *τ* ≥ 0 the 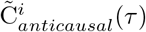 reveals the anticausal contribution (Figure 4c). This is how the results in Figure 4 were generated, where the causal (resp. anticausal) contribution is derived from the causal (resp. anticausal) model.

#### 4.7.1 Visualizing per region temporal contribution receptive field

Similar to the per region plot in Figure 3f, to generate a temporal contribution curve for each region (Figure 5), we average the spatial-temporal receptive field data (Figure 4) over to the same within-subject anatomical region labels. The control curve is generated by applying the same method for the shuffled model (grey curves in Figure 4). We omit those curves that are not significantly above noise level by Wilcoxon sign rank testing between averaged (over time) region contribution curves and the averaged (over time) noise level curve (see Extended Data Table 2).

## 5 Acknowledgements

We would like to thank Robert Knight and Sasha Devore for providing helpful comments on the manuscript. This work was supported by the National Science Foundation under Grant No. IIS-1912286 (Y.W. and A.F.) and National Institute of Health R01NS109367 (to A.F.).

## 6 Author contributions

R.W. conceived and implemented the decoding algorithm and interpreted the model with assistance from Y.W. and A.F.; X.C participated in the data processing and performance evaluation; A.K.G participated in the data processing and visualization; L.Y. participated in data acquisition, preprocessing, and visualization; P.D. and D.F. provided clinical care; W.D. provided neurosurgical clinical care; O.D. assisted in patient care and consent; Y.W. led the research project and advised from engineering perspective; A.F. co-led the project with Y.W., participated in all data acquisition, and advised from neuroscience perspective; R.W. and A.F. co-wrote the manuscript with input from all authors.

## A Extended Data

### A.1 Additional Decoding Framework Details

#### A.1.1 Differentiable speech synthesizer

In a traditional vocoder, speech is generated by switching between voiced and unvoiced content. Each content comes from an autoregressive system driven by a certain excitation signal that is either a harmonic signal or a white noise signal [11]. Inspired by such a process, we construct our speech synthesizer shown in Fig. 6. It consists of two pathways. The *voice pathway* generates a voiced component by driving a harmonic excitation with time-varying fundamental frequency (i.e., pitch) 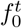 through a voice filter consisting of *N* formant filters, each described by a center frequency 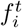 and an amplitude 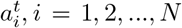. Note that we parameterize the bandwidth 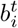 as a function of the center frequency 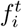, as discussed later. The *unvoice pathway* generates an unvoiced component by driving a white noise through an unvoice filter described as a center frequency 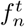, bandwidth 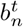 and amplitude 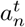 (in addition to the *N* formant filters for the voice pathway). These two components are adaptively combined with a time-varying mixing factor *α^t^*, controlling the relative contribution between voiced sounds (for sonorant phonemes including vowels and nasals) and unvoiced sounds (for voiceless plosives and fricatives such as /p/, /s/). The voiced plosives and fricatives (such as /b/, /z/) can be generated as a combination of voiced and unvoiced components. Finally, the combined signal is amplified by a loudness parameter *L_t_*. In our study, we used *N* = 6 formants. The synthesizer is driven by a total of 18 time-varying speech parameters, including the fundamental (or pitch) frequency 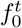, the mixing factor between the two pathways *α^t^*, the 12 parameters for the voice filter 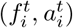 and the three parameters for the unvoice filter 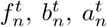, and the loudness *L^t^*. Given the parameter values at each time sample, the synthesizer can generate a spectrogram sample. The spectrogram is a differentiable function of the speech parameters so that we can back-propagate the gradient of the training loss in terms of the predicted spectrogram to the speech parameters, which can then be backpropagated to either the speech encoder or the ECoG decoder parameters. Specifically, let the *V^t^*(*f*) represent the spectrogram of the voicing component, *U^t^*(*f*) that of the unvoicing component, and *α^t^* ∈ [0, 1] the mixing factor. The combined spectrogram can be written as *S^t^*(*f*) = *α^t^V^t^*(*f*) + (1 – *α^t^*)*U^t^*(*f*). Finally, the synthesized speech spectrogram is 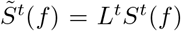, where *L^t^* is the loudness that modulates the signal cross time.

##### Formant filters in the voice pathway

The filter in the voice pathway consists of multiple formant filters, corresponding to the multiple formants associated with vowels. The formant filter shape over frequency, which is related to the resonance property of the vocal tract, is closely related to the timbre of speakers’ voice [32]. We have found that a predefined analytic form such as generalized Gaussian cannot cover all feasible filter shapes. Instead, we learn a speaker-dependent prototype filter for each formant based on the speaker’s natural speech. We represent the prototype filter (*G_i_*(*f*) for the *i*-th formant as a piecewise linear function, linearly interpolated from *g^i^*[*m*], *m* = 1…*M*, the amplitudes of the filter at *M* uniformly sampled frequencies up to *f_max_*. We restrict the resulting filter *G_i_*(*f*) to be unimodal (with a single peak of value 1) by properly constraining g[m]. Given *g*[*m*], *m* = 1…*M*, the peak frequency 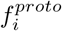 and the half-power bandwidth 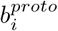 can be determined. The actual formant filter at any time can be written as a shifted and scaled version of *G_i_*(*f*). Specifically, at time *t*, given an amplitude 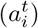, a center frequency 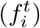, and a bandwidth 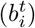, the *i*-th formant filter is given by

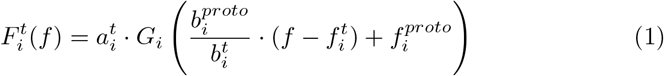

Then the filter for the voice pathway with *N* formant filters can be written as 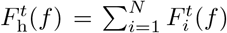. We learn the parameters *g*[*m*], *m* = 1…*M* for *G_i_*(*f*) during the unsupervised pre-training of the speech encoder, which does not require neural data. Fitting such a prototype filter is not data-hungry even with a relatively large *M*. We used *M* = 20 in our experiment. Although two formants (*N*=2) have been shown to suffice for intelligible reconstruction [7], we use *N*=6 in our experiments for more accurate synthesis. We denote the parameter set for the voice filter at time *t* by 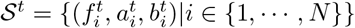. As explained later, the bandwidth 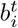 parameters are not independent speech parameters, rather functions of the center frequencies 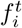.

##### Unvoice filter

For the unvoice pathway, we add a broadband filter described by 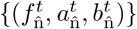. The shape of this filter 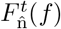 follows equation (1) but with the filter coefficients 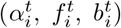 replaced by 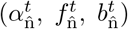. The bandwidth is constrained to satisfy 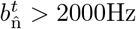, following the broadband nature of obstruent phonemes. We also keep the multiple formant filters in the voice filter described by *S^t^*. This is motivated by the fact that human beings differentiate consonants with similar sounds such as /p/ and /d/, not only by the immediate burst of these sounds, but also the development of the following formant frequency until the next vowel [33]. To encode such formant transitions, we use the same formant filter parameters for modeling the narrow passbands in both the voiced component and the unvoiced component. The parameter set for the unvoiced component is thus 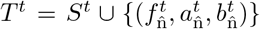. The overall filter for the unvoice pathway is: 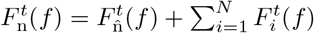.

To further reduce the parameter space dimension, we model the bandwidth 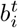 of a formant filter as a piecewise linear function of the center frequency 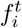. We assume

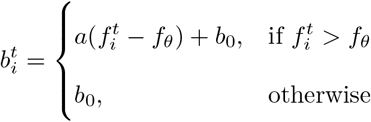

where threshold frequency *f_θ_*, slope *a*, and baseline bandwidth *b*_0_ are three parameters that can be learnt during unsupervised pre-training, shared among all formant filters.

##### Harmonic excitation

In the voice pathway, the voice filter is applied on the harmonic excitation. This pathway models the human production of vowels and nasals, which results from the voice excited by the vocal cord shaped by the vocal tract. The excitation is constructed by sinusoidal harmonic oscillations with a time varying fundamental frequency 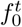. Inspired by the formulation in [12], we define the harmonic excitation *h^t^* as: 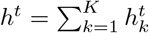, where *K* is the total number of harmonics ( K=80 in our experiment). Assuming the initial phase is 0, each harmonic resonance 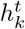 at time step *t* has an instant phase that is the accumulation of resonance frequency in the past. Specifically, the *k*-th resonance at time step *t* is 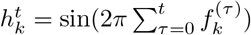, where 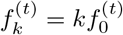. Denoting the spectrogram of *h^t^* as *H^t^*(*f*), the spectrogram of the voice component is the multiplication of *H^t^*(*f*) and the voice filter, i.e., 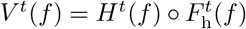.

##### Noise excitation

The unvoice pathway models consonants like plosives and fricatives, where the vocal tract and human mouth filter the airflow through the mouth. It follows a similar process as in the harmonic counterpart. The major difference is that the excitation being filtered becomes stationary white Gaussian distributed noise 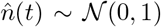, with a corresponding spectrogram *N^t^*(*f*). The filtered noise spectrogram (i.e., the unvoice component) is 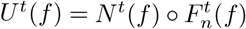.

#### A.1.2 ECoG decoder and speech encoder

The ECoG decoder is constructed by a three-dimensional ResNet that treats time-varying signals on an ECoG grid array as spatiotemporal three-dimensional tensors (width × height × time duration). As is depicted in Figure 6c, after an initial temporal convolutional layer (with 128 feature map filters and a kernel size of 1 × 1 × 9(72*ms*)), the signal passes through eight residual blocks. Each block contains two three-dimensional convolutional layers (with 128 feature map filters, each has kernel size of 3×3×5(40*ms*)). The output of the residual blocks creates a shared latent representation consisting of 128 feature maps (each is a one-dimensional temporal signal by average pooling the two spatial dimensions), which is then fed into different output heads (each applies each consists of one or two fully connected layers acting on the 128 features at the same time point) to generate speech parameters. The overall temporal receptive field for generating one speech parameter sample is 73 temporal samples of 584 ms.

The speech encoder network architecture we choose is as simple as possible to demonstrate the effectiveness of the speech synthesizer design. In the experiment, we use three layers of temporal convolution (we treat the frequency axis of the spectrogram as the feature dimension) to generate a latent representation (Figure 6d). Each convolutional layer has 128 feature maps and a temporal kernel size of 3 frames (24ms). To output the speech parameter, we apply the same multi-head structure to the latent representation as in the last layer of the ECoG decoder.

#### A.1.3 Loss and training hyper-parameters

The speech encoder is trained with a weighted average of the mixed spectral loss and the parameter loss. The mixed spectral loss [12] is defined as:

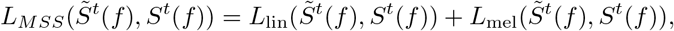

in which,

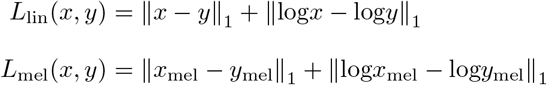

where *S^t^*(*f*) and 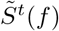 denote the ground truth and reconstructed spectrograms, respectively, subscript lin means that the frequency is in the linear scale while the subscript mel means the frequency is in the mel scale. In our experiments, we use 256 frequency samples (ranging from 0-8000 Hz) for both linear scale and mel scale speech sepctrograms.

Let’s denote the *j*-th reconstructed speech parameter as 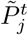 and its reference 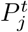, the overall training loss for the ECoG decoder becomes:

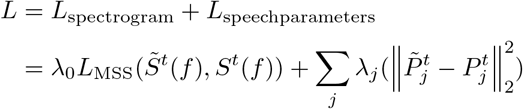

where λ*_j_* balance the contribution from different loss terms since they have different physical meanings and scales.

Both the speech encoder and ECoG decoder are fitted by Adam optimizer with hyper-parameters: *lr* = 10^-3^, *β*_1_ = 0.9, *β*_2_ = 0.999. We train an individual ECoG decoder and speech encoder per patient. The pre-training of the speech encoder and the training of the ECoG decoder share the same training/testing set partition.

## B Additional Figures and Tables

**Figure E1:**
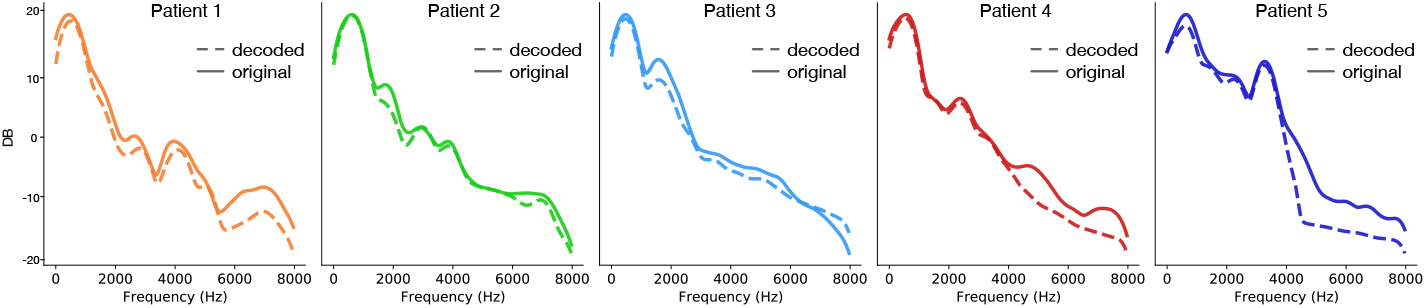
The spectral energy distribution of the decoded and original speech for five patients. Visualized by averaging the broadband spectrograms magnitude across time of all test samples.

**Figure E2:**
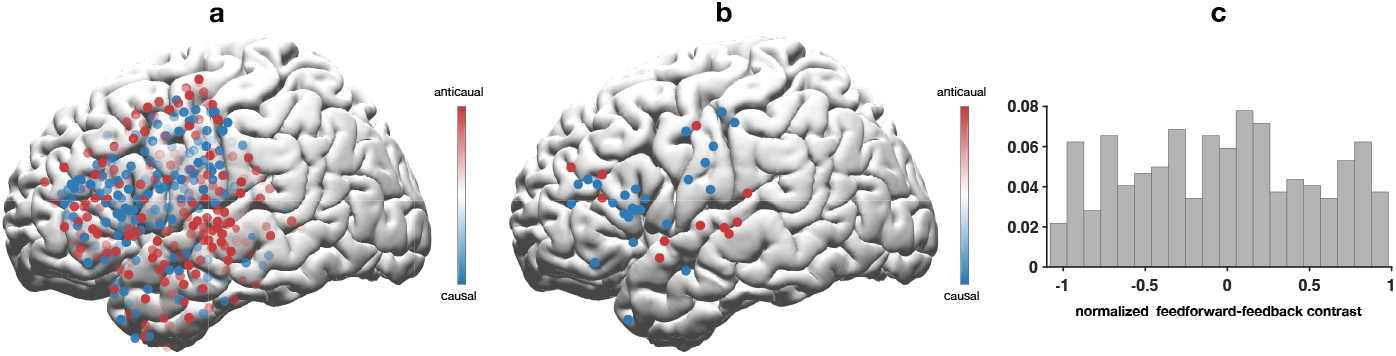
Normalized contrast of feedforward vs. feedback contribution. (a) electrode level feedforward-feedback contribution contrast, normalized by the sum of feedforward and feedback contribution magnitude. (b) electrodes with large feedforward-feedback polarity with the normalized contrast magnitude>0.9. (c) The histogram of the normalized contrast. Positive bins correspond to anticausal polarization.

**Figure E3:**
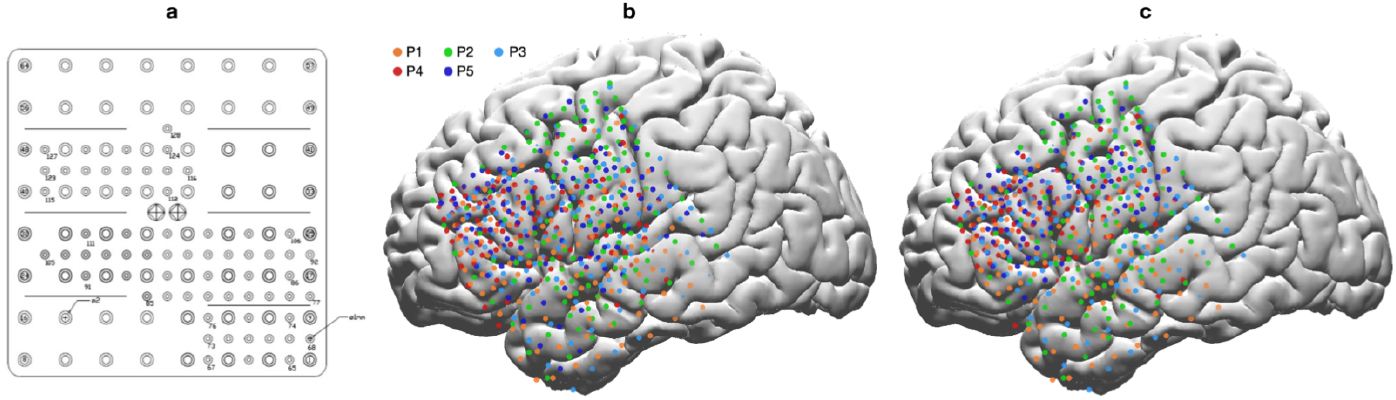
Electrodes array and implant location of all five patients (P1-P5) in our experiments. (a) The 128 electrodes on the hybrid density ECoG array. (b) All electrodes on cortex (MNI). (c) All electrodes with usable data. Only data from these electrodes are used to train the ECoG decoder models.

---

**Table 1:** Statistics of data in Figure 3f. The P-value and Z-value are reported for Wilcoxon sign rank test between feedback and feedforward contributions across all electrodes and test trials within each anatomical region. The Z-value represents the rank based test statistic with positive values reflecting anticausal contributions and negative values reflecting causal contributions.

| Anatomical region | p | z |
| --- | --- | --- |
| rSTG | 0.0332 | -2.9628 |
| cSTG | 1.607E-15 | 9.6234 |
| rMTG | 2.5108E-04 | 4.9359 |
| mMTG | 1.5257E-13 | 9.0185 |
| cMTG | 0.2269 | 1.5656 |
| ventralprecentral | 4.9511E-8 | -7.1409 |
| dorsalprecentral | 0.4349 | 0.6525 |
| postcentral | 6.419E-04 | -4.9612 |
| cMFG | 0.0248 | 3.1417 |
| rMFG | 0.1988 | 1.7202 |
| parstriangularis | 2.6715E-06 | 6.3518 |
| parsopercularis | 8.0693E-15 | -9.6185 |
| supramarginal | 1.1144E-04 | 5.3919 |

**Table 2:** Statistics of data in Figure 4 and 5. Per anatomical region P-value and Z-value are reported for Wilcoxon sign rank test between the each regions’ contribution and the shuffled model’s contribution (control curves in Figure 5. The Z-value represents the rank based test statistic with positive values reflecting larger real contributions compared with shuffled contributions. This is shown for the causal model (pre-production period), causal model (during production period), and anticausal model, respectively. Curves of each individual electrode and test trial are considered one sample, and are averaged across time to perform the Wilcoxon sign rank test. The red marked regions in the table are highlighted to show no significance (P-value>0.05) and are omitted when plotting the curves in Figure 5 as described (Method - Revealing delay-dependent contribution of different cortical regions from the trained ECoG to speech model - Visualizing per region temporal contribution receptive field).

| Anatomical region | Causal pre |  | Causal during |  | Anticausal |  |
| --- | --- | --- | --- | --- | --- | --- |
|  | p-value | z-value | p-value | z-value | p-value | z-value |
| cSTG | 0.7226 | 0.075 | 5.6745E-10 | 8.4078 | 1.2168E-22 | 10.9874 |
| rSTG | 0.4942 | -0.0510 | 1.2703E-07 | 6.8284 | 0.1557 | 1.9149 |
| mMTG | 5.3721E-06 | 6.2146 | 0.3689 | 0.2741 | 1.8216E-28 | 12.1658 |
| cMTG | 0.1671 | 1.0126 | 0.231 | -0.501 | 0.4781 | -0.3012 |
| rMTG | 5.1253E-19 | 10.1547 | 0.1293 | 2.1082 | 1.4923E-10 | 8.2051 |
| ventralprecentral | 1.7845E-58 | 16.2047 | 3.0286E-77 | 17.5451 | 2.2394E-60 | 17.1839 |
| dorsalprecentral | 2.9083E-12 | 8.0932 | 8.9452E-04 | 4.4590 | 1.4512E-09 | 7.9235 |
| postcentral | 3.67853E-91 | 21.4986 | 9.34051E-104 | 22.1393 | 6.9834E-34 | 14.0134 |
| supramarginal | 2.2905E-06 | 6.7810 | 0.5924 | -0.2945 | 1.8542E-07 | 6.9384 |
| parsopercularis | 3.9368E-76 | 19.0572 | 3.843E-72 | 18.5329 | 3.083E-04 | 5.3823 |
| parstriangularis | 7.2744E-77 | 19.5782 | 5.8573E-31 | 13.9374 | 2.0273E-37 | 14.4676 |
| rMFG | 2.3846E-27 | 12.2940 | 2.0371E-07 | 7.8460 | 0.3643 | 0.3823 |
| cMFG | 4.0274E-26 | 11.0042 | 2.83632E-07 | 6.9027 | 9.02834E-19 | 9.1881 |

**Table 3:** Statistics of data in Figure 4 and 5. Per anatomical region P-value and Z-value are reported for Wilcoxon sign rank test between the causal (during production period) model and the anticausal model (The positive/negative Z-values represent the direction of the contribution where positive values denote anticausal greater than causal), as well as the causal model between during- and pre-epochs (The positive/negative Z-values represent the direction of the contribution where positive values denote during production greater than pre-production). Curves of each individual electrode and test trial are considered as one sample, and are averaged across the time epoch to perform the Wilcoxon sign rank test. The red marked regions in the table are highlighted to denote no significance (P-value>0.05).

| Anatomical region | Causal vs. Anticausal |  | Causal during vs pre |  |
| --- | --- | --- | --- | --- |
|  | P-value | Z-value | P-value | Z-value |
| cSTG | 2.6789E-17 | 9.6711 | 4.718E-04 | 3.696 |
| rSTG | 0.0343 | -2.9457 | 6.2075E-04 | 4.7427 |
| mMTG | 3.2252E-13 | 9.0928 | 4.5863E-04 | -4.0475 |
| cMTG | 0.3930 | 1.0021 | 0.2718 | -1.1957 |
| rMTG | 1.8511E-04 | 5.1625 | 1.0173E-10 | -8.9283 |
| ventralprecentral | 2.8012E-15 | -10.0562 | 8.2757E-05 | 5.0475 |
| dorsalprecentral | 0.6492 | 0.2967 | 5.5615E-04 | -3.4394 |
| postcentral | 3.0581E-08 | -6.1286 | 0.3037 | 1.7462 |
| supramarginal | 1.9928E-07 | 6.0301 | 4.8257E-06 | -6.0274 |
| parsopercularis | 8.6228E-18 | -10.0274 | 0.5922 | 0.1582 |
| parstriangularis | 0.0162 | 3.9003 | 3.2532E-32 | -12.4583 |
| rMFG | 0.0021 | -4.9475 | 2.5714E-04 | -5.0131 |
| cMFG | 0.0045 | 3.9862 | 3.0747E-09 | -7.0652 |

**Table 4:** Peak time of each anatomical region curves in Figure 5 a,b,c. Each column reports the peak time of the temporal receptive field curves for the causal model (pre-production), causal model (during production), and anti-causal model, respectively. The peak of each region is calculated based on the averaged curve (averaged across trials and electrodes within the region).

| Anatomical region | Causal<br>(pre) | Causal<br>(during) | Anticausal |
| --- | --- | --- | --- |
| cSTG | - | -352 | 168 |
| rSTG | - | -256 | - |
| mMTG | -176 | - | 240 |
| cMTG | - | - | - |
| rMTG | -192 | - | 312 |
| ventralprecentral | -196 | -208 | 280 |
| dorsalprecentral | -192 | -184 | 144 |
| postcentral | -248 | -256 | 192 |
| supramarginal | -120 | - | 184 |
| parsopercularis | -248 | -280 | 232 |
| parstriangularis | -240 | -336 | 264 |
| rMFG | -248 | -304 | - |
| cMFG | -248 | -304 | 208 |

## References

[1] Hassan Akbari, Bahar Khalighinejad, Jose L Herrero, Ashesh D Mehta, and Nima Mesgarani. Towards reconstructing intelligible speech from the human auditory cortex. Scientific reports, 9(1):874, 2019.

[2] Miguel Angrick, Christian Herff, Garett Johnson, Jerry Shih, Dean Krusienski, and Tanja Schultz. Interpretation of convolutional neural networks for speech spectrogram regression from intracranial recordings. Neurocomputing, 342:145–151, 2019.

[3] Miguel Angrick, Christian Herff, Emily Mugler, Matthew C Tate, Marc W Slutzky, Dean J Krusienski, and Tanja Schultz. Speech synthesis from ecog using densely connected 3d convolutional neural networks. Journal of neural engineering, 16(3):036019, 2019.

[4] Gopala K Anumanchipalli, Josh Chartier, and Edward F Chang. Speech synthesis from neural decoding of spoken sentences. Nature, 568(7753):493–498, 2019.

[5] Kristofer E Bouchard, Nima Mesgarani, Keith Johnson, and Edward F Chang. Functional organization of human sensorimotor cortex for speech articulation. Nature, 495(7441):327–332, 2013.

[6] Edward F Chang, Caroline A Niziolek, Robert T Knight, Srikantan S Nagarajan, and John F Houde. Human cortical sensorimotor network underlying feedback control of vocal pitch. Proceedings of the National Academy of Sciences, 110(7):2653–2658, 2013.

[7] Edward F Chang, Kunal P Raygor, and Mitchel S Berger. Contemporary model of language organization: an overview for neurosurgeons. Journal of neurosurgery, 122(2):250–261, 2015.

[8] Josh Chartier, Gopala K Anumanchipalli, Keith Johnson, and Edward F Chang. Encoding of articulatory kinematic trajectories in human speech sensorimotor cortex. Neuron, 98(5):1042–1054, 2018.

[9] Connie Cheung, Liberty S Hamilton, Keith Johnson, and Edward F Chang. The auditory representation of speech sounds in human motor cortex. Elife, 5:e12577, 2016.

[10] Trinity B Crapse and Marc A Sommer. Corollary discharge across the animal kingdom. Nature Reviews Neuroscience, 9(8):587–600, 2008.

[11] Li Deng and Douglas O’Shaughnessy. Speech processing: a dynamic and optimization-oriented approach. CRC Press, 2018.

[12] Jesse Engel, Lamtharn Hantrakul, Chenjie Gu, and Adam Roberts. DDSP: Differentiable digital signal processing. arXiv preprint arXiv:2001.04643, 2020.

[13] James L Flanagan. Speech analysis synthesis and perception, volume 3. Springer Science & Business Media, 2013.

[14] Mario Fleischer, Silke Pinkert, Willy Mattheus, Alexander Mainka, and Dirk Mürbe. Formant frequencies and bandwidths of the vocal tract transfer function are affected by the mechanical impedance of the vocal tract wall. Biomechanics and modeling in mechanobiology, 14(4):719–733, 2015.

[15] Adeen Flinker, Edward F Chang, Heidi E Kirsch, Nicholas M Barbaro, Nathan E Crone, and Robert T Knight. Single-trial speech suppression of auditory cortex activity in humans. Journal of Neuroscience, 30(49):16643–16650, 2010.

[16] Adeen Flinker, Anna Korzeniewska, Avgusta Y Shestyuk, Piotr J Franaszczuk, Nina F Dronkers, Robert T Knight, and Nathan E Crone. Redefining the role of broca’s area in speech. Proceedings of the National Academy of Sciences, 112(9):2871–2875, 2015.

[17] Joaquin M Fuster. The prefrontal cortex—an update: time is of the essence. Neuron, 30(2):319–333, 2001.

[18] Joaquin M Fuster. Upper processing stages of the perception–action cycle. Trends in cognitive sciences, 8(4):143–145, 2004.

[19] Joaquín M Fuster. The prefrontal cortex in the neurology clinic. Handbook of clinical neurology, 163:3–15, 2019.

[20] Jeremy DW Greenlee, Roozbeh Behroozmand, Charles R Larson, Adam W Jackson, Fangxiang Chen, Daniel R Hansen, Hiroyuki Oya, Hiroto Kawasaki, and Matthew A Howard III. Sensory-motor interactions for vocal pitch monitoring in non-primary human auditory cortex. PloS one, 8(4):e60783, 2013.

[21] Jeremy DW Greenlee, Adam W Jackson, Fangxiang Chen, Charles R Larson, Hiroyuki Oya, Hiroto Kawasaki, Haiming Chen, and Matthew A Howard III. Human auditory cortical activation during self-vocalization. PloS one, 6(3):e14744, 2011.

[22] Frank H Guenther. A neural network model of speech acquisition and motor equivalent speech production. Biological cybernetics, 72(1):43–53, 1994.

[23] Frank H Guenther. Neural control of speech. Mit Press, 2016.

[24] Kaiming He, Xiangyu Zhang, Shaoqing Ren, and Jian Sun. Deep residual learning for image recognition. In Proceedings of the IEEE conference on computer vision and pattern recognition, pages 770–778, 2016.

[25] Christian Herff, Lorenz Diener, Miguel Angrick, Emily Mugler, Matthew C Tate, Matthew A Goldrick, Dean J Krusienski, Marc W Slutzky, and Tanja Schultz. Generating natural, intelligible speech from brain activity in motor, premotor, and inferior frontal cortices. Frontiers in neuroscience, 13:1267, 2019.

[26] Gregory Hickok. Computational neuroanatomy of speech production. Nature reviews neuroscience, 13(2):135–145, 2012.

[27] Gregory Hickok. The cortical organization of speech processing: Feed-back control and predictive coding the context of a dual-stream model. Journal of Communication Disorders, 45(6):393–402, 2012. 21st Annual NIDCD-Sponsored ASHA Research Symposium (2011):Neuroplasticity in the Mature Brain.

[28] Gregory Hickok. The architecture of speech production and the role of the phoneme in speech processing. Language, Cognition and Neuroscience, 29(1):2–20, 2014.

[29] Gregory Hickok and David Poeppel. The cortical organization of speech processing. Nature Reviews Neuroscience, 8(5):393, 2007.

[30] John F Houde and Srikantan S Nagarajan. Speech production as state feedback control. Frontiers in human neuroscience, 5:82, 2011.

[31] Colin Humphries, Merav Sabri, Kimberly Lewis, and Einat Liebenthal. Hierarchical organization of speech perception in human auditory cortex. Frontiers in neuroscience, 8:406, 2014.

[32] Eric J Hunter, Jan G Švec, and Ingo R Titze. Comparison of the produced and perceived voice range profiles in untrained and trained classical singers. Journal of Voice, 20(4):513–526, 2006.

[33] Jintao Jiang, Marcia Chen, and Abeer Alwan. On the perception of voicing in syllable-initial plosives in noise. The Journal of the Acoustical Society of America, 119(2):1092–1105, 2006.

[34] Eric R Kandel, James H Schwartz, Thomas M Jessell, Steven Siegelbaum, A James Hudspeth, and Sarah Mack. Principles of neural science, volume 4. McGraw-hill New York, 2000.

[35] John Kominek, Tanja Schultz, and Alan W Black. Synthesizer voice quality of new languages calibrated with mean mel cepstral distortion. In Spoken Languages Technologies for Under-Resourced Languages, 2008.

[36] Sergey Korolev, Amir Safiullin, Mikhail Belyaev, and Yulia Dodonova. Residual and plain convolutional neural networks for 3d brain mri classification. In 2017 IEEE 14th international symposium on biomedical imaging (ISBI 2017), pages 835–838. IEEE, 2017.

[37] Joseph G Makin, David A Moses, and Edward F Chang. Machine translation of cortical activity to text with an encoder–decoder framework. Nature Neuroscience, 23(4):575–582, 2020.

[38] Brian T Miller and Mark D’Esposito. Searching for “the top” in top-down control. Neuron, 48(4):535–538, 2005.

[39] Muge Ozker, Werner Doyle, Orrin Devinsky, and Adeen Flinker. Cortical network underlying speech production during delayed auditory feedback. bioRxiv, 2021.

[40] Ramprasaath R Selvaraju, Michael Cogswell, Abhishek Das, Ramakrishna Vedantam, Devi Parikh, and Dhruv Batra. Grad-cam: Visual explanations from deep networks via gradient-based localization. In Proceedings of the IEEE international conference on computer vision, pages 618–626, 2017.

[41] Jennifer Shum, Lora Fanda, Patricia Dugan, Werner K Doyle, Orrin Devinsky, and Adeen Flinker. Neural correlates of sign language production revealed by electrocorticography. Neurology, 95(21):e2880–e2889, 2020.

[42] Kristina Simonyan, Hermann Ackermann, Edward F Chang, and Jeremy D Greenlee. New developments in understanding the complexity of human speech production. Journal of Neuroscience, 36(45):11440–11448, 2016.

[43] Daniel Smilkov, Nikhil Thorat, Been Kim, Fernanda Viégas, and Martin Wattenberg. Smoothgrad: removing noise by adding noise. arXiv preprint arXiv:1706.03825, 2017.

[44] Donald T Stuss and Robert T Knight. Principles of frontal lobe function. Oxford University Press, 2013.

[45] Cees H Taal, Richard C Hendriks, Richard Heusdens, and Jesper Jensen. A short-time objective intelligibility measure for time-frequency weighted noisy speech. In 2010 IEEE international conference on acoustics, speech and signal processing, pages 4214–4217. IEEE, 2010.

[46] Ran Wang, Xupeng Chen, Amirhossein Khalilian-Gourtani, Zhaoxi Chen, Leyao Yu, Adeen Flinker, and Yao Wang. Stimulus speech decoding from human cortex with generative adversarial network transfer learning. In 2020 IEEE 17th International Symposium on Biomedical Imaging (ISBI), pages 390–394. IEEE, 2020.

[47] Ran Wang, Yao Wang, and Adeen Flinker. Reconstructing speech stimuli from human auditory cortex activity using a WaveNet approach. In 2018 IEEE Signal Processing in Medicine and Biology Symposium (SPMB), pages 1–6. IEEE, 2018.

[48] Andrew I Yang, Xiuyuan Wang, Werner K Doyle, Eric Halgren, Chad Carlson, Thomas L Belcher, Sydney S Cash, Orrin Devinsky, and Thomas Thesen. Localization of dense intracranial electrode arrays using magnetic resonance imaging. Neuroimage, 63(1):157–165, 2012.

[49] James F.A. Poulet and Berthold Hedwig. The cellular basis of a corollary discharge. Science, 311: 518–522, 2006.

[50] Eliades, Steven J., and Xiaoqin Wang. Neural substrates of vocalization feedback monitoring in primate auditory cortex. Nature, 453(7198):1102–1106, 2008.

